# Comparative Analysis of DNA Structural Parameters and the Corresponding Computational Tools to Differentiate Regulatory DNA Motifs and Promoters

**DOI:** 10.1101/2024.03.26.584893

**Authors:** Vasumathi Manivelan, Basavaraju Kavyashree, Bindu Sadanandan, Sravanti Vaidya, Kshitish K Acharya

## Abstract

Analyzing and distinguishing functionally distinct DNA regions is crucial for various applications, including predicting DNA motifs and promoters, and exploring the mechanisms of gene expression regulation in disease conditions. Our understanding of mammalian promoters, particularly those associated with differentially expressed genes (DEGs), particularly remains limited. However, existing methods for such analysis require refinement. Despite the value of DNA Structural Parameters (DSPs), users often struggle to objectively select parameters and tools, especially given the limited options available. This study addresses this challenge by thoroughly investigating DSP-tool combinations – particularly the local structural parameters that can be analyzed via web-interfaces, with a goal to discern human DNA motifs and promoters. What sets this study apart are the following aspects: a) examination of disease-associated promoters; b) attention to regulatory specific DNA motifs; c) compilation and comparison of all publicly available online tools and parameters for analyzing DNA structures, and test all available DSP-tool combinations. Through the execution of over half a million queries, the study identified DSP-tool combinations that consistently outperformed others in differentiating DNA sequences across various types of analyses. Notably, the ‘propeller twist’ emerged as a standout DSP, while DNAshape, complemented by DNAshapeR scripts, demonstrated exceptional performance among the tools across four distinct types of analyses: testing motifs, sequences post motif insertion, comparing promoters with control sequences, and analyzing promoters of genes either up- or down-regulated under disease conditions. Significant alterations were observed in the values of multiple DSPs for 100-nucleotide-long promoter and control sequences following the insertion of single motifs such as triplex target sites (TTS), quadruplex-forming structural motifs (QSFM), and transcription factor binding sites (TFBS). However, no single combination proved universally successful. For instance, when analyzing promoters associated with differential gene expression, different combinations proved effective across various human diseases. These findings provide valuable insights for making informed selections among available options for sequence-based queries.

## Introduction

Understanding the structural aspects of nucleic acids is paramount as they play a pivotal role in a wide spectrum of molecular interactions (1,2). These interactions, in turn, serve as the groundwork for a plethora of molecular processes, spanning from transcription initiation (3) to chromosomal recombinations (4), chromatin interactions, and RNA splicing (5,6). In the domain of transcription regulation, particularly within motifs and promoters, the significance of DNA structure is profound. Numerous researchers have meticulously scrutinized DNA structural features associated with promoters and their diverse types. It is widely acknowledged that the structural characteristics of prokaryotic promoters starkly differ from those of eukaryotic promoters (7). Notably, in bacterial promoter regions proximal to the Transcription Start Site (TSS), traits such as stability, rigidity, and curvature exhibit notable distinctions (8,9). Efforts to predict promoters based on DNA structural features have been abundant (10,11). However, it is crucial to acknowledge the motif-based diversity inherent among promoters when delving into structural analysis. Consideration of such diversity is pivotal for a comprehensive understanding of the implications of DNA structure on transcription regulation.

In eukaryotic species with intricate body organization, such as mammals, the differential expression of genes is a well-established phenomenon essential for development and maintenance across various tissues and stages (1,2). Beyond development, genes exhibit distinct expression patterns within each tissue or cell type, under different environmental and physiological conditions, including during disease states. Among the myriad of regulatory mechanisms, transcription initiation emerges as a pivotal phase for gene expression control. The diversity in promoters plays a crucial role in governing the initiation of transcription, a process central to gene expression regulation. Grouping promoters based on the expression patterns of associated genes proves to be a valuable approach. Notably, promoters linked to different gene expression patterns often display distinct distributions of DNA motifs (12–14).

In the complex landscape of eukaryotic promoter regions, a multitude of B-DNA motifs, including Transcription Factor Binding Sites (TFBSs) (15–17), as well as non-B-DNA motifs such as Triplex Target Sites (TTSs) (18) and Quadruplex Structure-Forming Motifs (QSFMs) (19–21), are anticipated to play integral roles. These motifs are intricately associated with genes that exhibit differential expression patterns (22,23). In contrast to bacterial promoters, eukaryotic counterparts are often characterized by a significantly higher degree of complexity and diversity (24,25), owing to the diverse nature of DNA motifs (26). Notably, within the genome of Saccharomyces cerevisiae, most Transcription Factors (TFs) seem to harbor at least one conserved DNA structural parameter within their binding sites (16). Furthermore, investigations into eukaryotic genomes have revealed distinct structural features between promoters with and without TATA motifs across six eukaryotic genomes (27).

The exploration of structural aspects within promoters has been somewhat limited, particularly concerning Triplex Target Sites (TTSs) and Quadruplex Structure-Forming Motifs (QSFMs). TTSs, characterized by polypurine stretches, hold potential significance in gene expression regulation. For instance, in the ETS2 promoter, TTSs positioned approximately 40 base pairs upstream of the Transcription Start Site (TSS) have been identified to modulate gene expression dynamics by impeding SP1 binding (28). On the other hand, QSFMs, comprised of guanine- or cytosine-rich DNA sequences, have garnered attention. Studies have revealed an over-representation of G-quadruplex motifs in promoters associated with genes exhibiting high expression levels and nuclease hypersensitivity (29).

The observations made thus far suggest a promising avenue wherein the potentially diverse structural features of motifs could serve as discriminators between promoters and random sequences, or even among promoters associated with different gene expression patterns. Despite these promising indications, such possibilities remain largely unexplored, especially within complex genomes like the human genome. To bridge this gap, computational tools have been developed to facilitate the analysis of DNA sequences, offering invaluable assistance to researchers across various domains. Many of these tools rely on local structural parameters to delineate DNA characteristics. For instance, DNAshape stands out as a user-friendly tool with a web interface, enabling biologists to analyze sequences effectively. However, it currently provides analysis based on only four DNA structural parameters (DSPs).

While such tools present researchers with accessible means to embark on structure-based characterization of diverse DNA regions, including functionally distinct human promoters, a critical need arises to compile a comprehensive list of these tools and evaluate their efficacy in discerning key DNA regions. The potential variability in the lengths of DNA motifs, especially certain Transcription Factor Binding Sites (TFBSs), underscores the necessity of considering multiple local or dinucleotide structural parameters. While numerous studies on promoters have traditionally emphasized global structural features like DNA bendability or stability, it’s crucial to acknowledge that these global characteristics are often shaped by dinucleotide parameters.

However, the intricate nuances of local DNA structural parameters (DSPs) in the assessment of higher eukaryotes demand attention. Many user-friendly tools, tailored for researchers with limited programming skills, specifically exploit local DSPs. This prompts a deeper exploration into their effectiveness in discerning various types of promoters within species characterized by complex body organization, such as mammals. Such investigations carry significant potential for accelerating our understanding of the mechanisms governing differential gene expression, particularly in disease states. Hence, bridging the current emphasis on global structural parameters with the yet-to-be-explored potential of local DSPs, notably those available in existing tools, is pivotal for unraveling the intricacies of eukaryotic gene regulation.

When engaging with computational programs, users are confronted with the challenge of selecting from a plethora of DNA Structural Parameters (DSPs) (30) to effectively analyze DNA sequences. However, there currently exists no established scientific basis, particularly from the user’s perspective, for choosing specific computational tools or DSPs tailored to analyze DNA motifs or longer promoter sequences. Despite the recognition of this gap, and concerted efforts to systematically compare tools (31,32), such studies have faced several limitations, leaving users to rely on subjective choices. Hence, there is a pressing need for a comprehensive listing and systematic comparison of suitable DSPs.

In this study, we meticulously compiled and compared DNA Structural Parameters (DSPs) along with the currently available tools for DNA structure analysis. We rigorously tested various DSP-tool combinations to assess their efficacy in differentiating: (a) short DNA motifs from each other, (b) promoter or other DNA sequences based on the presence or absence of specific DNA motifs, (c) promoters versus random sequences, and (d) functionally diverse promoter sets within a species. Furthermore, we conducted a comparative analysis to gauge the relative efficiency of DSP-tool combinations across these specific types of analyses.

## 2. Methods

In our endeavor to evaluate the effectiveness of different combinations of computational tools and DNA Structural Parameters (DSPs), we embarked on a meticulous process. Initially, we meticulously compiled all available options. Subsequently, we conducted a total of 589,720 distinct queries aimed at distinguishing between various types of DNA sequences. a) Short (8-12 nucleotides) DNA motifs: To assess DSP-tool combinations, we scrutinized variations in DSP values across 100 motifs. b) Promoters and control sequences with and without the insertion of 70 DNA motifs: Our objective was to pinpoint DSP-tool combinations showcasing statistically significant differences in values post-insertion of specific motifs into 100 nucleotide-long sequences. We meticulously compared values across 50 promoter and 50 control sequences before and after motif insertion. The 70 tested motifs encompassed ten Triplex Target Site motifs (TTSs), ten Quadruplex Forming Structural Motifs (QSFMs), and ten Transcription Factor Binding Sites (TFBSs) from each of five TF superclasses. c) Promoters (∼100 or 300 nucleotides each) vs. control sequences: We delved into the values obtained using DSP-tool combinations for 500 core and extended promoters, juxtaposed against an equal number of control sequences. d) Functionally distinct (850 nucleotides long) promoters: Our analysis involved a comparison of values obtained using DSP-tool combinations for human promoters associated with opposing gene expression patterns, both under normal and disease conditions. The specifics of each comparison are meticulously outlined below, shedding light on the intricacies of our evaluation process.

### 2.1 Listing the computational tools and DNA Structural Parameters (DSPs) for analysing DNA sequences

To compile research articles pertinent to our investigation, we focused on several key topics: computational tools developed for DNA structure analysis, DNA Structural Parameters (DSPs), and previous comparative studies in these domains. We initiated a comprehensive literature search employing carefully curated search terms tailored to each topic. While PubMed served as our primary resource due to its superior precision, we also leveraged a couple of other search engines to ensure comprehensiveness (33). Our quest led us through a meticulous screening process of relevant research reports and reviews, culminating in the compilation of a comprehensive list of tools adept at analyzing DNA structures. Each potential tool underwent rigorous scrutiny through multiple queries using random inputs, enabling us to evaluate their functionalities from a user’s perspective. Some tools were excluded from further consideration due to functional limitations. Subsequently, among the tools that successfully underwent queries with DNA sequences, we short-listed those deemed suitable for comparative studies. Meanwhile, the compilation of DNA Structural Parameters (DSPs) was conducted following an exhaustive literature review, supplemented by screening DiProDB, a notable resource in the field (34).

For the current comparative study, we identified and short-listed four tools: cgDNA (35), DNAshape (36), DNAshapeR (37), and Pse-in-One 2.0 (38). Concurrently, fifteen DNA Structural Parameters (DSPs) were chosen for detailed analysis. Within the cgDNA framework, 12 DSPs were deemed relevant for querying DNA sequences via the Octave platform (39). On the other hand, the online version of DNAshape was capable of processing structural values for only four DSPs. In contrast, the DNAshapeR package demonstrated a more robust capability, accommodating 14 DSPs through the backend of DNAshape. Consequently, we leveraged the intermediate output of the DNAshapeR package to obtain values for these 14 DSPs from DNAshape. Further-more, DNAshapeR was employed independently to derive 14 DSP values, furnishing normalized DSP values using the prior-DSP values of DNAshape. In the case of Pse-in-One, six inter-base pair DSPs were considered. During querying, we employed the pseudo dinucleotide composition (PseDNC) method with lambda (sequence order information) set to 0, along with a weight factor parameter of 0.1.

### 2.2 Identifying and analyzing short DNA motifs

In the study, a total of 100 motifs, varying in length from 8 to 12 nucleotides, were employed. Among these motifs, 22 were recognized for their involvement in DNA-protein complexes, while 33 were sourced from the Nucleic Acid Database (NDB) (40), representing typical B-DNA motifs. Notably, a considerable proportion of these motifs displayed a GC-bias. Consequently, to ensure representation across nucleotide compositions, an additional set of 45 motifs enriched with AT-rich elements was generated.

Subsequently, DSP values were estimated for all motifs utilizing the four selected tools. The mean absolute DSP values were computed for each motif, following which the Coefficient of Variation (CV), calculated as the standard deviation divided by the mean, was determined across motifs for each DSP.

### 2.3 Identifying promoter sequence sets for analysis

The determination of human genes expressed ubiquitously across tissues involved a meticulous process. Initially, a set of ubiquitous human genes, as identified by a study employing microarray-based expression data across multiple tissues (41), underwent further refinement. These genes were short-listed based on their expression across normal adult tissues, leveraging mammalian gene expression databases: MGEx-Tdb (42), MGEx-Udb (43), and MGEx-Kdb (unpublished). The reliability score obtained from these databases gauged the consistency of gene transcription across similar conditions but different datasets/experiments. Cumulative reliability scores from the three databases facilitated the hierarchical arrangement and selection of ubiquitous genes, with the top 500 genes considered. Additionally, ubiquitous genes from the microarray datasets were short-listed if they exhibited high reliability scores across the three tissues under consideration.

Similarly, testis-specific genes were initially obtained from the TiGER database. Those testis-specific genes from TiGER, which were also transcribed according to MGEx-Tdb, were short-listed, provided they displayed a dormant status in uterus and kidney tissues, as indicated by MGEx-Udb and MGEx-Kdb. The reliability scores (scaled 0–10) from MGEx-Tdb and the EST enrichment scores (scaled 0–10) were aggregated to establish a hierarchical list of tissue-specific genes, with the top 200 testis-specific genes considered. To facilitate further analysis, the RefSeq IDs of the ubiquitous and testis-specific genes were converted to Ensembl IDs of transcripts using ‘BioMart’ (https://www.ensembl.org/info/data/biomart/index.html) for promoter extraction. Additionally, the study considered 416 up-regulated genes and 354 down-regulated genes in Non-Obstructive Azoospermia (NOA).

### 2.4 Comparing sequences before and after insertion of individual motifs

The analysis focused on the −250 to −150 promoter regions of the top 50 ubiquitous genes. Given that core promoters, typically situated between +50 to −100 nucleotides, are often enriched with transcription factor binding sites (TFBSs), we opted for slightly distant promoter regions as test sequences. To create controls, random versions of promoter sequences were generated using the shuffle function in the List::Util Perl module, based on the top 50 promoter sequences.

For motif insertion, a variety of elements were utilized. This included three Triplex Target Sites (TTSs) sourced from the Nucleic Acid Database (NDB) (40), along with seven artificially generated TTSs spanning lengths from 10 to 17 nucleotides. Additionally, ten distinct Quadruplex Forming Structural Motifs (QSFMs) from NDB (40) were employed. Furthermore, ten Transcription Factor Binding Sites (TFBSs) were selected from each of the following transcription factor superclasses: the basic domain, immunoglobulin fold, zinc-coordinating binding domain, helix-turn-helix domain, and other all-alpha helical binding domains. Consensus sequences for these TFBSs were procured from the TFClass database (47). In total, 70 motifs of three main types—TFBSs, TTSs, and QSFMs—were individually inserted into the middle of each promoter or control sequence, with an equal number of existing nucleotides being replaced.

Subsequently, DNA Structural Parameter (DSP) values were estimated for each sequence from the two sets (promoter and control) both before and after the insertion of these 70 motifs, one motif at a time. The mean of absolute DSP values was computed for each sequence using the selected tools. Henceforth, the ‘means of DSP values of the sequences’ in a set were referred to as ‘Sequence-Set-Mean’ for each DSP. The paired Student t-test was then employed to compare Sequence-Set-Means for each set before vs. after the motif insertion.

### 2.5 Comparing core and larger promoter regions with control sequences

The analysis encompassed the promoter sequences of the top 500 ubiquitous genes, as outlined previously, alongside their corresponding control sequences. To generate control sequences, the top 500 promoter sequences underwent random shuffling across positions using the shuffle function within the List::Util Perl module. Both core promoter regions, spanning from −100 to +1, and larger regions between −250 and +50, along with their corresponding control sequence sets, were incorporated into the study.

Sequence-Set-Means were computed across promoter and control sets, and subsequently compared using an unpaired t-test, as detailed earlier. Additionally, the average value for each DNA Structural Parameter (DSP) for each position within a set was calculated. This average of position-specific DSP-values for the entire set is henceforth denoted as ‘Position-Mean’ for each DSP.

To assess the distribution of Position-Mean within the promoter set compared to the control set, a two-sample Kolmogorov-Smirnov (KS) test was conducted, following the methodology described by Massey in 1951 (48).

### 2.6 Comparing function-based promoter sets

Promoter regions (−750 to +100 regions, with +1 representing the Transcription Start Site) were meticulously extracted utilizing an in-house script for both the top 200 ubiquitous and top 200 testis-specific transcripts, as previously delineated. Concurrently, akin promoter regions for up- and down-regulated genes in Non-Obstructive Azoospermia (NOA) were retrieved from the Eukaryotic Promoter Database (EPD) (49).

The DNA Structural Parameter (DSP) values of these promoters were meticulously estimated leveraging cgDNA, DNAshape, and DNAshapeR. Notably, Pse-in-One was excluded from this analysis due to its inability to provide position-wise DSP values. Subsequently, the Position-Mean across the sequences was meticulously calculated for every promoter and control set, mirroring the methodologies employed in previous analyses. Following the computation of Position-Mean DSP values for each promoter set, comparisons were drawn against other promoter sets and the corresponding control sequences. The disparity was quantified utilizing the two-sample Kolmo-gorov-Smirnov (KS) test on the Position-Mean of promoter and control sets.

### 2.7 Querying the tools with sequences, for specific DSPs

The initiation of queries for each tool was carried out manually, one at a time. Due to the necessity for multiple queries for each tool, scripts were developed to streamline the process. Specifically, scripts were crafted in MATLAB for cgDNA, Python for Pse-in-One, and R for both DNAshape and DNAshapeR. To ensure the accuracy and reliability of the automated scripts, they underwent validation by comparing their outputs with those obtained from manual queries. For added confirmation, the online version of DNAshape was employed in a few rounds of validation.

The comprehensive analysis encompassed a total of 4,600 automated DNA motif queries, each contributing to the calculation of coefficients of variation (CVs). In the realm of DSP-value assessment in promoters before and after motif insertion, a substantial 322,000 queries were executed. Furthermore, an additional 92,000 queries were performed to compare promoter regions of two different lengths with control sequences. Another facet of the study involved 171,120 queries conducted for comparing function-based promoter sets.

Scripts were written to submit multiple queries in cgDNA and to utilize DNAshape for extracting more DSPs via the DNAshapeR module.

## 3. Results

### 3.1 DNA Structural Parameters (DSPs) are compiled for comparative analysis

Among the extensive array of dinucleotide properties cataloged in DiProDB, a total of 65 conformational and physicochemical parameters (DSPs) pertinent to DNA were meticulously selected for analysis. These encompassed a broad spectrum of characteristics, including global helical parameters such as ‘tip’ and ‘inclination’, inter-base-pair parameters like ‘shift’, groove-related parameters such as ‘width’ and ‘distance’, and energy-related parameters like ‘entropy’. However, properties specific to RNA and those reliant on letter-based criteria were excluded from further consideration to maintain focus.

In addition to the parameters sourced from DiProDB, literature was thoroughly reviewed to identify other potential DSPs. This compilation yielded a comprehensive set of properties including DNA bending stiffness, protein-induced deformability, DNA denaturation, duplex disrupt energy, hydroxyl radical cleavage pattern, and two tri-nucleotide-based properties—DNaseI and nucleosome position preference. Furthermore, to enhance the scope of analysis, additional parameters such as electrostatic potential (EP), Helical Twist (HelT), and intra-base-pair parameters like ‘buckle’, ‘opening’, ‘stretch’, ‘stagger’, and ‘shear’ were incorporated from parameter lists utilized in certain tools.

The culmination of this rigorous selection process resulted in a final set comprising 78 distinct DSPs deemed suitable for analyzing DNA regions. However, it is worth noting that only 15 parameters are currently under consideration by the four tools compatible with sequence-based queries, as elucidated below.

### 3.2 Useful computational tools for examining DNA structures are listed, and some of them selected for comparative analysis

There exist 16 tools designed to estimate specific values of DNA structural properties, as detailed at https://startbioinfo.org/cgi-bin/prelimresources.pl?tn=DNA-RNA%20structures for additional information). These tools exhibit variations in methodology: some determine values for each pair of nucleotides within a given DNA structure from scratch, while others calculate values for each query sequence based on predefined sets of values (prior values) per parameter for nucleotide pairs (across the strand) or dinucleotides (neighbors on a strand).

Tools accepting DNA structures in PDB format as queries employ the de novo method, whereas those accepting DNA sequences as queries utilize prior values. Considering that the majority of DNA regions across all species lack established structures, our study prioritized sequence-based prediction tools—specifically DNAshape, cgDNA, DNAshapeR, and Pse-in-One—for further investigation. It is important to note that the selected tools employ distinct methods and prior values for estimating DSP values.

### 3.3 Useful DSP-tool combinations are identified for distinguishing short patterns (motifs)

When queried with each motif, the tools consistently generated diverse values for every DNA Structural Parameter (DSP). The primary objective was to discern the optimal combination of tools and DSPs capable of distinguishing between motifs effectively. Therefore, the Coefficient of Variation (CV) of means of DSP-values across 100 motifs was computed for each parameter using each of the four tools, and subsequently compared (Table 1). Notably, Pse-in-One failed to exhibit variations in mean-values across queried motifs.

**Table 1.** Coefficient of Variation for the means of 100 motifs.

| Parameters | Tools & CV values of means of motifs |  |  |  |
| --- | --- | --- | --- | --- |
|  | cgDNA | DNASHape | DNASHapeR | Pse-in-one |
| Buckle | 0.3491 | 0.5416 | 0.091 | 0 |
| Opening | 0.295 | 0.537 | 0.137 | 0 |
| Propeller-twist | 0.1472 | 0.383 | 0.25 | 0 |
| Shear | 0.271 | 0.251 | 0.173 | 0 |
| Stagger | 0.228 | 0.392 | 0.144 | 0 |
| Stretch | 0.185 | 0.231 | 0.315 | 0 |
| Shift | 0.345 | 0.34 | 0.047 | 0.0009 |
| Slide | 0.186 | 0.0724 | 0.267 | 0.00096 |
| Rise | 0.0081 | 0.0083 | 0.096 | 0.0009 |
| Roll | 0.158 | 0.279 | 0.125 | 0.0009 |
| Tilt | 0.446 | 0.441 | 0.04 | 0.00096 |
| Twist | 0.0195 | 0 | 0 | 0.0009 |
| Helical-twist | 0 | 0.0156 | 0.1683 | 0 |
| MGW | 0 | 0.0521 | 0.115 | 0 |
| EP | 0 | 0.142 | 0.125 | 0 |

Utilizing cgDNA, DNAshape, or DNAshapeR, a CV of ≥ 0.2 was observed across motifs for multiple DSPs. The DSP-tool combinations demonstrating the most promising CV included ‘buckle’, ‘opening’, ‘tilt’, ‘stagger’, and ‘propeller twist’ with DNAshape, alongside ‘tilt’ with cgDNA. In total, 18 out of the 46 DSPs-tool combinations exhibited potential for effectively differentiating short DNA motifs.

### 3.4 Useful DSP-tool combinations are identified for distinguishing 100-nt-long DNA sequences based on their motif contents

The presence of Transcription Factor Binding Sites (TFBSs) and other specific motifs defines the characteristics of promoter sequences, potentially inducing identifiable structural alterations in DNA sequences. This hypothesis was rigorously tested by inserting a total of 70 motifs, one at a time, into each 100-nucleotide DNA sequence. These motifs comprised Triplex Target Sites (TTSs) (10 motifs), Quadruplex Structure-Forming Motifs (QSFMs) (10 motifs), and TFBSs (50 motifs), with ten TFBSs allocated to each specific superclass of Transcription Factors (TFs).

The trends observed in the variations among the DNA Structural Parameter (DSP) values for these 70 motifs closely mirrored the findings of the earlier analysis involving 100 motifs. DSP-values were collected for each sequence both before and after motif insertion, and a mean value was computed for each set of promoter sequences per DSP per tool. These Sequence-Set-Means of DSPs for 50 promoter sequences were then compared before and after each motif insertion. Similarly, values for 50 control sequences were compared before and after motif insertions.

While Pse-in-One did not yield significant differences in any DSP-values of sequence-sets before and after motif-insertions, several other DSP-tool combinations exhibited notable DSP-value differences after TFBS insertions. Particularly, properties that displayed significant differences in at least 8 out of 10 motif-insertion cases for each TF-superclass were identified (Table 2). For instance, the parameter ‘shear’ emerged as effective in differentiating motif-insertions using cgDNA or DNAshape across four TF-Superclasses.

**Table 2.** Common DNA Structural Parameters (DSPs) across tools that seem to work for differentiating 100nt long sequences before vs. after inserting TFBSs of different superclasses of TFs (TF-superclasses)

| **Domains of TF-super-classes corresponding to the TFBSs inserted in promoter sequences** | **Common significant properties across tools** | **Tools** |
| --- | --- | --- |
| Basic DNA binding domain | Shift | cgDNA, DNashape |
|  | MGW | DNashape, DNashapeR |
| Zinc-coordinating binding domain | Shift, Shear, Stretch | cgDNA, DNashape |
| Helix-turn helix domain | Opening, Propeller twist, Roll, Rise, Shear | cgDNA, DNashape, DNashapeR |
|  | Stretch, Rise | cgDNA, DNashape |
|  | Slide, Stagger | DNashape, DNashapeR |
| Other alpha-helical | Opening, Propeller twist, Rise, Shear, Roll, Stagger | cgDNA, DNashape, DNashapeR |
|  | Tilt | cgDNA, DNashape |
|  | Slide, Helical twist | DNashape, DNashapeR |
| Immunoglobulin domain | Propeller twist, Shear | cgDNA, DNashape |
|  | Stagger | cgDNA, DNashapeR |
|  | Helical twist, Rise, EP | DNashape, DNashapeR |

Conversely, fewer DSPs exhibited significantly different values in ≥ 8 of 10 cases (motif-insertions) with other tools. Notably, no differences were observed between promoter sequences as a set compared to the control sequences, regardless of motif-insertions. When using cgDNA, DNAshape or DNAshapeR, ‘propeller twist’, ‘roll’ and ‘tilt’ values were different after inserting TTSs, while values for ‘propeller twist’, ‘slide’ ‘rise’, ‘shear’, ‘twist’ and ‘helical twist’ changed after QSFM-insertions. Many other DSP-tool combinations also showed significant differences after TTSs and QSFMs insertions (Table 3). Interestingly, no differences were observed between promoter sequences as a set compared to the control sequences – with or without the motifinsertions.

**Table 3.** DNA Structural Parameters (DSPs) across tools that seem to work with 100% success rate for differentiating 100nt long sequences before vs. after inserting TFBSs of different superclasses of TFs (TF-superclasses)

| Domains of TF-superclasses corresponding to the TFBSs inserted in promoter sequences | Significant DSP with 100% success rate for differentiating the TFBSs | Tools |
| --- | --- | --- |
| Basic DNA binding domain | Shift | cgDNA |
| Zinc-coordinating binding domain | Buckle | cgDNA |
|  | Stretch | DNashape |
| Helix-turn helix domain | EP, propeller twist, roll, Shear | DNashape |
| Other alpha-helical | Stagger | DNashape, DNashapeR |
|  | Buckle | DNashapeR |
| Immunoglobulin domain | Shear | cgDNA |
|  | Rise | DNashape |

Overall, among the 46 DSP-tool combinations, only specific combinations proved effective in differentiating DNA sequences after insertion of TFBSs corresponding to each TF-superclass. More-over, certain combinations could discern sequences after TTS or QSFM insertions, further emphasizing the nuanced impact of motifs on DNA structural parameters.

### 3.5 Useful DSP-tool combinations are identified for differentiating promoter regions from random sequences

The tools employed in the analysis utilize distinct prior dinucleotide values or algorithms to estimate each DNA Structural Parameter (DSP) value in the query sequences. Since the effectiveness of each DSP and tool in differentiating promoter sequences was not previously compared, this analysis was undertaken. Both smaller (core; −100 to +1) and larger (proximal; −250 to +50) promoter regions were tested using various DSP-tool combinations, and the resulting DSP-values were compared with those obtained from the corresponding control sequences.

The Sequence-Set-Mean value for each DSP was compared between each promoter and control set. Notably, no significant differences were observed in mean DSP-values across sets when using Pse-in-One. Conversely, with the other three tools, many DSP-values differed significantly (p-value <0.05) between the promoter and control sequences. These combinations included ‘stretch’, ‘twist’, and ‘roll’ in cgDNA, ‘MGW’, ‘propeller twist’, ‘slide’, ‘stretch’, ‘tilt’, ‘stagger’, and ‘roll’ in DNAshape, and ‘roll’, ‘MGW’, ‘slide’, and ‘propeller twist’ in DNAshapeR.

Highly significant differences in values for ‘stretch’ in cgDNA (P=0), ‘MGW’ in DNAshape (P=0), and DNAshapeR (P=0), and ‘roll’ in DNAshapeR (P=0) were observed for both the core and larger promoter sequences compared to the control sequence set. ‘Twist’ of cgDNA (P<7.5E-11) could also reliably distinguish core as well as larger promoter sequences from the corresponding control sequences. Many other DSP-tool combinations could reliably distinguish promoters from control sequences, with some combinations providing remarkably different values (P<0.0001) between the promoter and control sequences (Table 4).

**Table 4:**
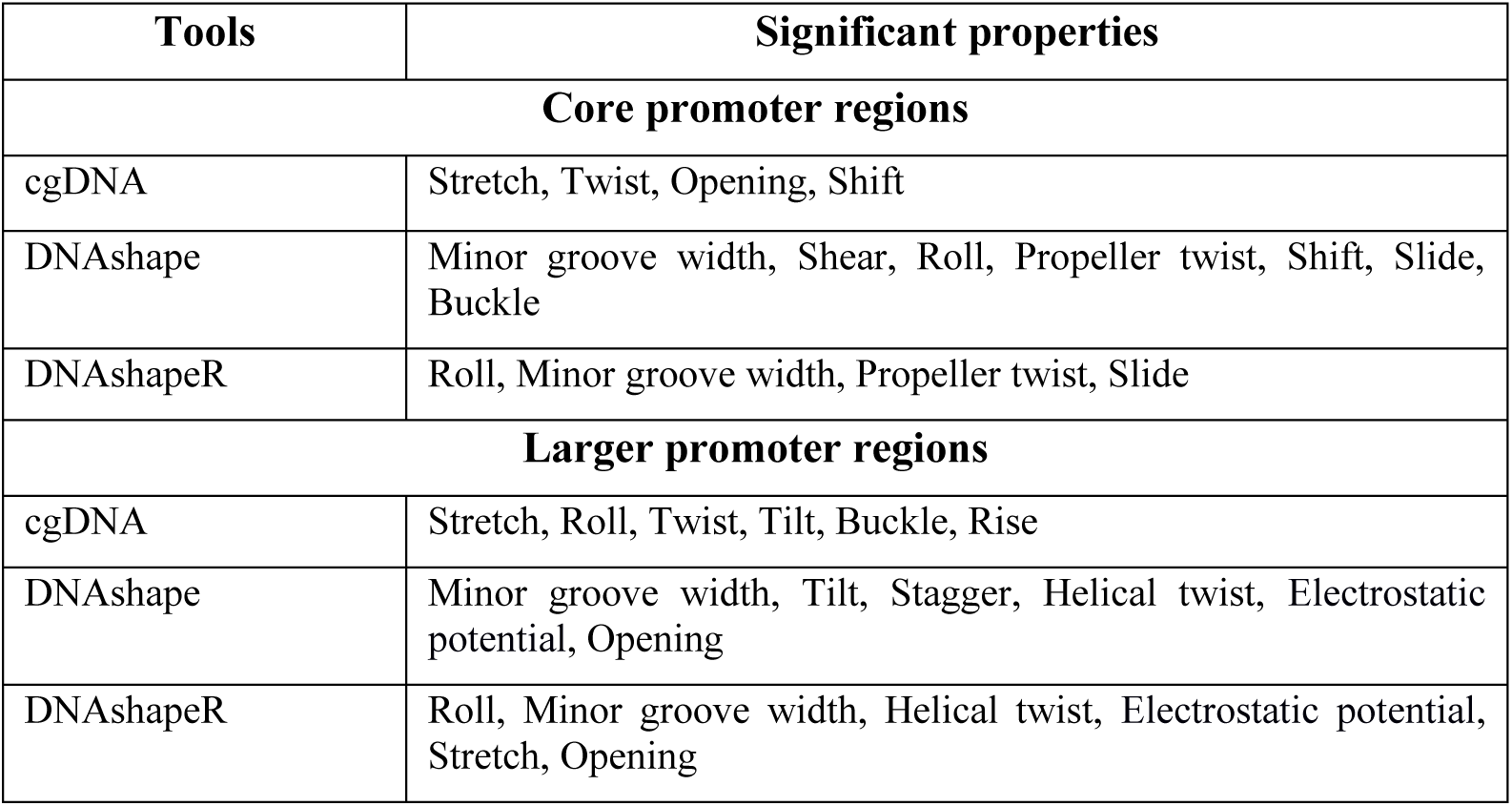
DNA Structural Parameters (DSPs) for which significantly different values (p<0.0001) were produced with promoter sequences compared to the control sequences.

Graphical representations of the Position-Mean values across the length of promoters revealed subtle differences in the patterns of these values across positions (Figures 1A & 1B). However, comparison of the overall values, Sequence-Set-Means, may not reveal obvious differences. To address this, cumulative-fractions (Kolmogorov-Smirnov or KS plot) of Position-Means from the promoter and control query sequences were plotted for each DSP and tool combination. Comparing the central values from such KS-plots provided a more straightforward assessment of the overall difference across the sequence sets.

**Figure 1.**
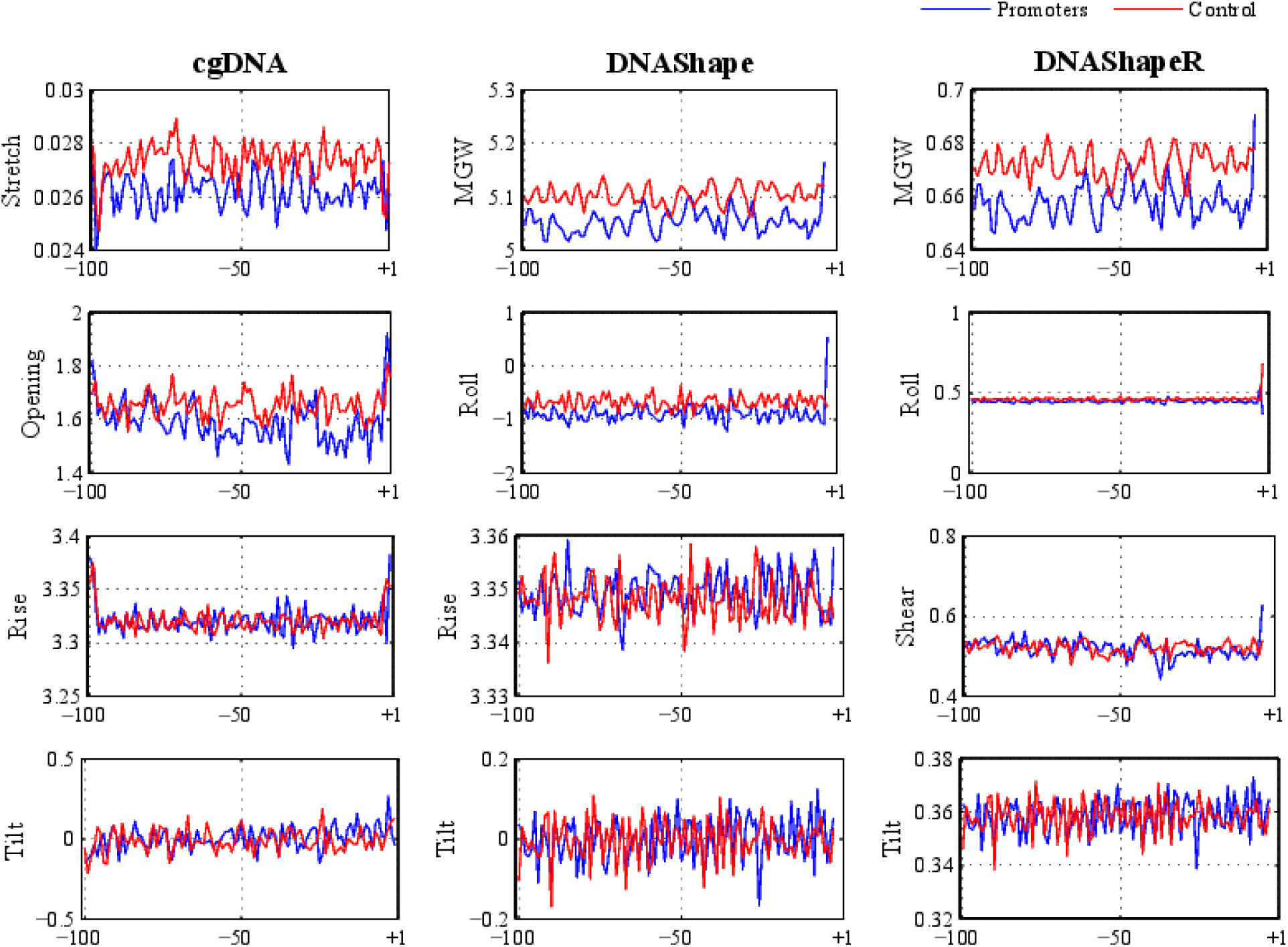
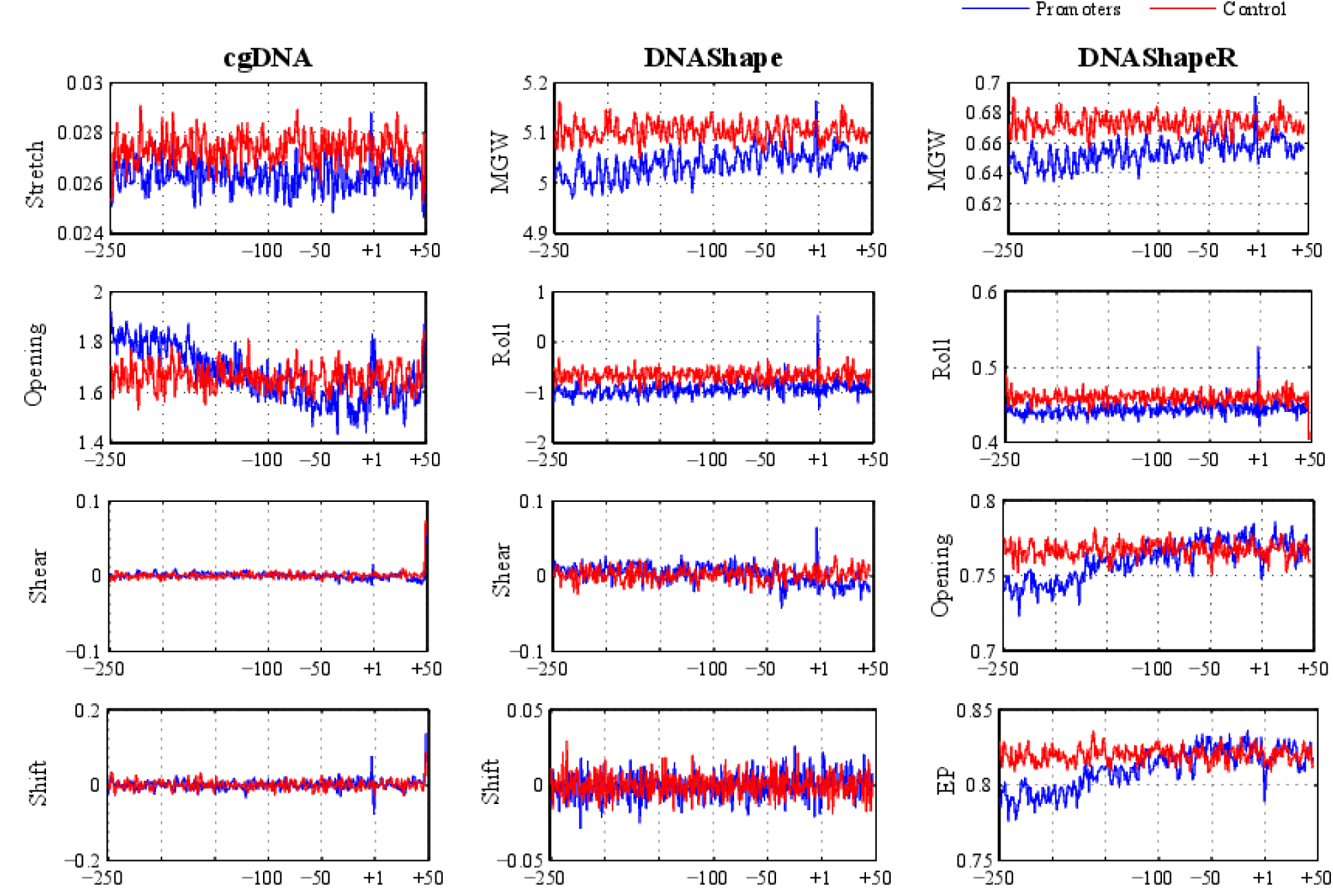
Position-Mean of DNA Structural Parameters (DSPs) at each position across 500 core promot-ers (blue) and an equal number of the control (randomly shuffled) sequences (red). The nucleotide posi-tions shown on the x-axis are with respect to the Transcription Start Site (TSS: +1) of the promoter se-quence. The tool name is displayed at the top of each vertical panel, along with the DSPs displayed in the y-axis. 1A. Results of a few DSP-tool combinations for smaller promoter regions (100 nucleotide length) 1B. Results of a few DSP-tool combinations for larger regions (300 nucleotides length).

The analysis identified statistically significant (P<0.05) differences for five more combinations when using KS-plots. These new combinations include ‘stagger’ and ‘propeller twist’ of cgDNA, ‘EP’ of DNAshape, and ‘buckle’, ‘opening’, and ‘stagger’ of DNAshapeR. With cgDNA, DNAshape, and DNAshapeR tools, ‘propeller twist’, ‘opening’, and ‘stagger’ combinations were useful for distinguishing core and large promoters from the controls. Overall, 20 of the 46 DSP-tool combinations could reliably differentiate core and large promoters from the controls.

### 3.6 Useful DSP-tool combinations are identified for differentiating function-based promoter-sets

In complex eukaryotes like mammals, promoters serve as pivotal elements in the regulation of gene expression, contributing to the diversity of gene functions. Thus, promoters associated with specific functional categories may exhibit distinct structural characteristics. We sought to investigate whether DNA structural features could effectively distinguish such functionally diverse sets of promoters using various DSP-tool combinations. Following a similar approach to previous tests, Position-Means within each function-based cluster were computed using cgDNA, DNAshape, and DNAshapeR tools. Corresponding values were also derived for the control sequences. We examined promoters linked to different gene expression patterns: ubiquitous expression, tissue-specific expression, and regulation in Non-Obstructive Azoospermia (NOA).

Our analysis revealed notable changes in DSP-values across the promoter sets, particularly around the proximal promoter regions (positions −150 to +100). Through KS tests comparing the respective controls, we identified consistent combinations of tools and parameters that yielded distinct results for promoters across sub-types. These combinations encompassed ‘stretch’, ‘stagger’, ‘opening’, ‘propeller twist’, ‘twist’, and ‘roll’ in cgDNA, and ‘MGW’, ‘roll’, ‘EP’, ‘opening’, ‘stretch’, ‘propeller twist’, ‘slide’, and ‘helical twist’ in DNAshape (Figures 2) and DNAshapeR.

**Figure 2.**
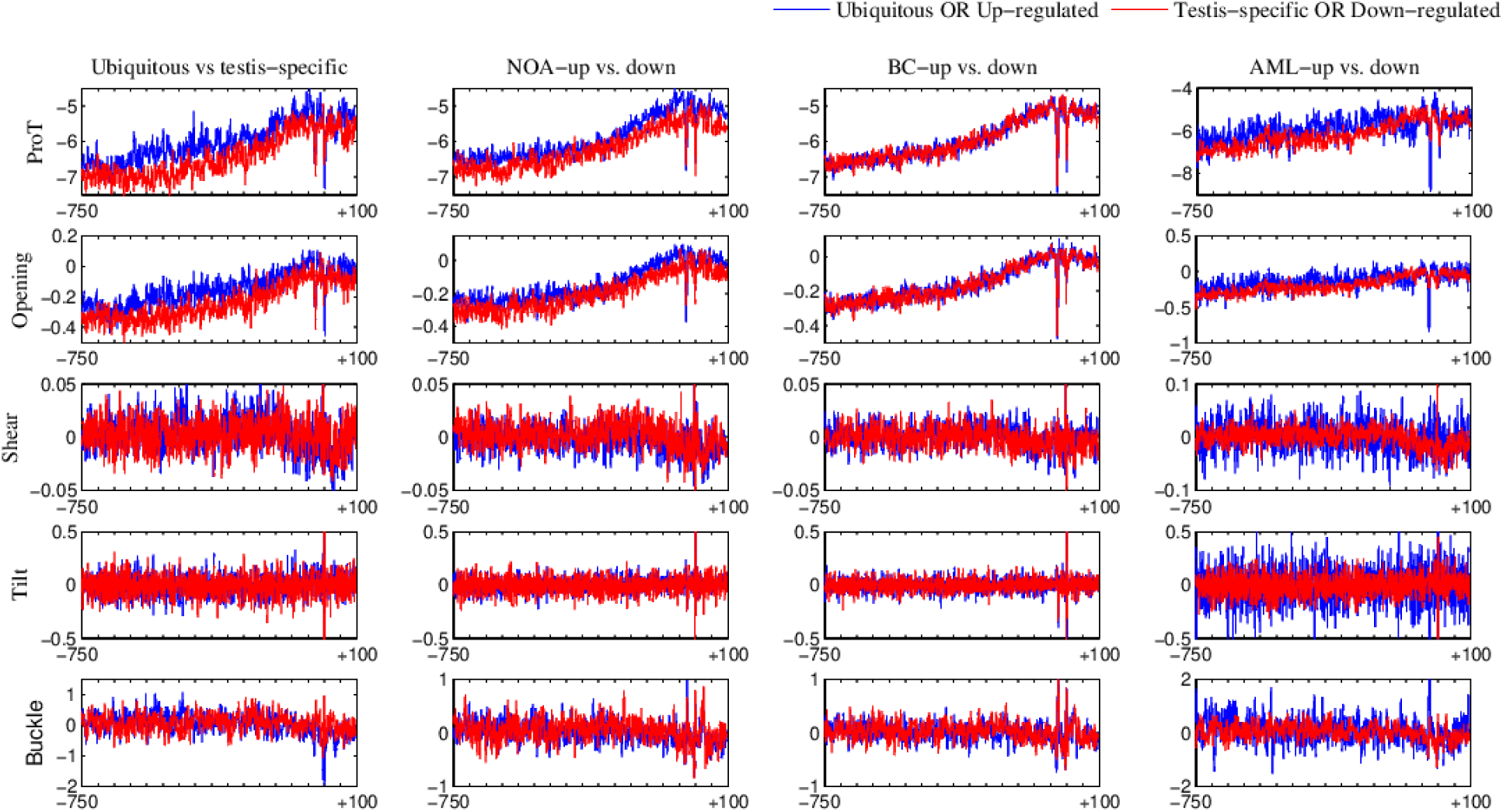
Position-Means of a few DNA Structural Parameters (DSPs) from DNAshape for differentiation of ubiquitous (blue line) vs. tissue-specific (red line) promoters and up (blue)- and down (red)-regulated genes promoters in non-obstructive azoospermia (NOA), breast cancer (BC) and acute myeloid leukemia (AML). The nucleotide positions shown are with respect to the transcription start site (TSS).

Promoters associated with divergent functional categories exhibited varying Position-Mean values for numerous DSPs throughout the promoter length. Specific properties such as ‘opening’, ‘propeller twist’, ‘roll’, and ‘stagger’ in cgDNA, and ‘propeller twist’, ‘EP’, ‘opening’, ‘slide’, ‘stretch’, ‘helical twist’, ‘stagger’, and ‘MGW’ in DNAshape and DNAshapeR facilitated differentiation between promoter subtypes such as ubiquitous vs. testis-specific, NOA up vs. down regulated. ‘Propeller twist’, ‘opening’, and ‘stagger’ with cgDNA, DNAshape, or DNAshapeR proved effective in differentiating most functionally varying promoters. It’s worth noting that analyzing the distribution of position-means alone may not directly reveal significant differences between two sets.

In summary, 22 out of 40 DSP-tool combinations reliably differentiated each promoter set from the control sequences, with 18 of these combinations also effectively discriminating functionally varying promoters in three comparisons. Ongoing research indicates that is possible to similarly discriminate promoters of other disease-associated genes, such as the genes differentially expressed under certain cancer conditions.

### 3.7 A comparison of results across 14 types of DNA-analyses

The analysis revealed that Pse-in-One did not provide useful results. However, ‘propeller twist’ in combination with DNAshape proved to be the most frequently effective DSP. Interestingly, this DSP also demonstrated high utility when used with cgDNA and DNAshapeR. Moreover, ‘roll’, ‘stagger’, and ‘opening’ showed promise in differentiating promoter sequences across many instances, particularly when paired with cgDNA, DNAshape, or DNAshapeR. Combining multiple DSPs with different tools may further enhance the success rate. For instance, the combination of ‘propeller twist’ and ‘shear’ showed promising results in distinguishing most promoter sequences when utilizing DNAshape.

Although no single combination proved universally effective, certain DSP-tool combinations emerged as reliable options for distinguishing a variety of DNA sequences. Additionally, it’s worth noting that specific promoter sets exhibited better distinguishability with higher DSP-tool combinations compared to others. Conversely, genes up- and down-regulated in other conditions provided varying results, with ‘shear’ being the only DSP showing consistent differentiation.

## 4. Discussion

The structural analysis of DNA holds promise in elucidating correlations between structural features and promoter function, potentially enabling the prediction of promoter activity. Such endeavors could significantly impact the exploration of disease-associated molecular mechanisms, given the well-documented variations in transcriptional profiles under pathological conditions. However, the procedural aspects of DNA analysis have been ambiguous, necessitating a comprehensive examination of the structural characteristics of DNA motifs and promoters. It’s crucial to evaluate the efficiency of procedural options, including available tools and structural parameters suitable for various types of analyses.

There is a noticeable dearth of comparative studies focusing on the tools and parameters utilized in DNA structure analysis, particularly from a user’s perspective. Previous studies in this domain primarily concentrated on DNA motifs and utilized tools capable of processing DNA structures as queries. However, tools utilizing sequence inputs have not been systematically compared for analyzing different types of DNA structures. The lack of comprehensive and systematic comparative studies on bioinformatic resources in general may stem from the time-consuming nature of such endeavors, as observed in studies related to protein-protein interaction databases, literature search tools, and gene expression databases. As highlighted in some of these reports, it is impractical for researchers with broader biological research objectives to undertake detailed comparative analyses of bioinformatic resources independently. Therefore, dedicated research efforts are necessary to thoroughly compare computational tools, databases, and parameters available for specific biological analyses, particularly from a user’s perspective. This includes comprehensive evaluations of promoter analysis based on structural parameters.

The current study provides a comprehensive overview of DNA Structural Parameters (DSPs) and their variants, along with the computational tools available for analyzing DNA structures. A total of 78 distinct structural properties were identified based on their physicochemical and conformational types, catering to the analysis of DNA molecules. However, existing sequence-based query-permitting tools utilize only a subset of these DSPs, predominantly focusing on local DSPs, which encompass inter- and intra-base-pair properties. These local DSPs serve as the building blocks for understanding global DNA structural properties such as ‘bendability’ and ‘curvature’. For instance, ‘wedge’ measures variations in ‘roll’ and ‘tilt’, while ‘curvature’ represents the cumulative effect of local dinucleotide wedge deflections along the DNA molecule. Moreover, variations in ‘slide’ impact parameters like ‘minor groove width’ and ‘minor groove depth’. Some available tools, such as DNAshape and DNAshapeR, facilitate the analysis of such parameters critical for shape-dependent recognition by DNA-binding proteins.

With four tools available for DNA sequence analysis, each considering multiple DSPs — up to 14 in a single tool — users can explore 46 DSP-tool combinations. In the current study, analyses were conducted in four phases following the listing of DSP-tool combinations. The initial phase aimed to evaluate the potential of all combinations in distinguishing short motifs. Previous studies have suggested that transcription factor binding sites (TFBSs) exhibit greater flexibility in human promoter regions compared to other genomic regions. Despite limited variation within each TFBS, there is considerable diversity in nucleotide sequences across DNA motifs that function as TFBSs, making them likely to possess distinguishable DSP values. While specific DSPs have been tested in some cases, such as the conservation of the ‘roll’ parameter in certain TFBSs, this study represents the first comprehensive analysis of multiple DNA motifs, particularly with various DSP-tool combinations. The results suggest that certain combinations can distinguish DNA motifs more effectively than others.

In the second phase of our analysis, we delved into whether any DSP-tool combinations could effectively differentiate between 100nt-long sequences based on the presence or absence of various types of motifs, including superclasses of transcription factor binding sites (TFBSs), Triplex Target Sites (TTSs), and Quadruplex Forming Structural Motifs (QSFMs) in the sequences. Previous studies have examined certain DSPs of eukaryotic promoters within the context of common core promoter elements. For instance, Yella and Bansal (2017) compared promoters with and without the TATA box, observing differences in structural features such as DNA stability, bendability, and curvature. Similarly, Vanaja and Yella (2022) compared promoters in the context of additional elements like Inr, BRE-u, and BRE-d, finding that promoters with certain regulatory motifs exhibit distinct structural properties. However, our study marks the first investigation into the effect of the presence or absence of a large number (70) of TFBSs in sequences using various combinations of tools and DSPs.

Several new insights have emerged from this study. For instance, in many cases, the values for ‘shear’ and ‘propeller twist’ changed remarkably upon the insertion of a single TFBS in a promoter sequence using DNAshape and cgDNA. The findings suggest that no single DSP-tool combination can reliably distinguish every type of DNA motif or sequence. However, specific DSP-tool combinations may prove valuable in multiple types of analyses. For instance, combinations involving ‘propeller twist’ and tools like DNAshape, cgDNA, or DNAshapeR appear effective in many test cases comparing sequences with and without individual motifs, including TTSs, QSFMs, and 30 TFBSs across TF-superclasses. This combination also reliably distinguishes promoters from control sequences and shows promise in differentiating functionally distinct promoter sets from each other.

Studies on the analysis of functionally distinct promoters have been limited. Kumar and Bansal (2017) demonstrated that DNA structural features vary among the promoters of highly expressed versus lowly expressed prokaryotic genes. To the best of our knowledge, the current results represent the first instance where: a) Structural properties of human or any other mammalian promoters are shown to vary depending on their association with distinct gene expression patterns across conditions. b) Specific combinations of currently available DSPs and tools have been identified that can differentiate functionally distinct eukaryotic promoters.

Even within each of the four types of comparisons performed in the current study, no combination appears 100% reliable. However, in the second type of comparison, reliable combinations were revealed for each subset—specifically, when studying the influence of the insertion of specific types of motifs. The parameter ‘tilt’ in cgDNA, DNAshape, and DNAshapeR tools exhibited a 100% success rate for distinguishing sequences after Triplex Target Site (TTS) insertion. Similarly, the ‘slide’ parameter in all the tested tools showed an almost 100% success rate for Quadruplex Forming Structural Motifs (QSFMs) in the sequences. Additionally, combinations involving DNAshape and parameters such as ‘electrostatic potential’ (EP), ‘propeller twist’, ‘roll’, and ‘shear’ showed a similar success rate in differentiating DNA sequences after the insertion of Transcription Factor Binding Sites (TFBSs) corresponding to the helix-turn-helix domain superclass.

The analysis had to be limited to DSP-tool combinations currently available for DNA sequence queries. While the results indicate that some of these combinations can effectively differentiate DNA motifs and promoters in many cases, there is scope for an even more comprehensive comparison. For example, there are indications that other structural parameters could aid in distinguishing promoters. The EP3 program, for instance, applies the base-stacking property, while PromPredict utilizes DNA duplex-free energy. Developing tools that enable users to test DNA sequences with additional parameters is desirable. Similarly, there is a need to explore more types of nucleic acid sequences, such as chromosomal recombination sites, and to test a wider range of promoter types with varying lengths. Furthermore, it is crucial to investigate the use of additional DSPs and their combinations.

Given the diversity in methods used across tools, it is unsurprising that the tools produced different results for identical query DNA sequences, even while considering identical DSPs. In the current study, a few DSPs from DNAshape and DNAshapeR displayed varying results for sequences following motif insertions, despite these tools utilizing the same prior values for their estimation of final DSP values. Earlier studies also demonstrated variations in results when using tools capable of predicting structural values for specific parameters using DNA structures in PDB format. Nevertheless, the current study design overlooked methodological differences and instead focused on testing DSP-tool combinations for specific applications. Such a user’s perspective is crucial concerning the final application of the tools. Studies conducted along similar lines are likely to identify one or more permutations of DSPs that can reliably distinguish promoter features and other types of functionally significant nucleic acid regions.

## 5. Conclusions

Overall, while none of the DSP-tool combinations proved universally reliable for all types of DNA regulatory region analyses, our study revealed specific cases where certain combinations demonstrated reliability, albeit to varying degrees. These findings offer valuable insights into streamlining the selection process among existing DSP-tool combinations for structure-based differentiation of DNA sequences. Moreover, they illuminate the inherent limitations in current DSP-tool combinations for analyzing structural features of motifs, promoters, or other DNA regions. For example, one limitation highlighted by our results is the inability of available tools to utilize all DSPs. This underscores the necessity for the development of new or enhanced versions of existing tools capable of considering a broader range of DSPs beyond the current 15. Additionally, these tools should employ multiple methods for estimating DSP values to enhance accuracy and comprehensiveness. Furthermore, the additional scripts and procedural guides provided in our study can greatly assist users in analyzing various DNA sequence sets of interest, facilitating more efficient and effective research in this field. Our results reveal that DSP values vary within promoter or other sequences of 100 nt length due to the presence of a single DNA motif, such as different TFBSs, TTSs, and QSFMs. Moreover, our findings represent the first instance where specific combinations of DSP-tools have been identified to distinguish between human promoters associated with distinct gene expression patterns. This discovery holds significant promise for exploring molecular mechanisms associated with cancer and other diseases, potentially opening new avenues for targeted research and therapeutic interventions.

## Author Contributions

KA conceptualized the study and planned it. VM performed most of the work, including the resource compilation, their comparison and the final comparative analysis, data summarization and visualization. KB obtained the clusters of differentially expressed genes. VM, KA, BS and SV contributed to the manuscript preparation.

## Acknowledgements

Dr. S. Thiyagarajan, IBAB, Bengaluru, India, identified limitations in some of the earlier data visualization methods. Dr. Sravanti Davuluri, Shodhaka Life Sciences Private Limited (https://www.shodhaka.net/), Bengaluru, helped us to identify key differentially expressed gene lists in cancer context. The research facility at IBAB is supported by the Department of IT, BT and S&T, Government of Karnataka, India.

